# UELer: a Jupyter-based framework for interactive exploration of multiplexed imaging datasets

**DOI:** 10.64898/2026.08.21.745700

**Authors:** Yu-Le Wu, Chun-Shan Liu, Bénédicte Lenoir, Kilian Merz, Michael T. Dill, Felix J. Hartmann

## Abstract

**Summary:** Multiplexed imaging and spatial proteomics generate complex datasets that require both computational analysis and visual inspection. However, these tasks mostly occur in separate environments because interactive viewers generally require a local display or an additional data server beyond the remote Jupyter sessions itself where large datasets are computationally analyzed. We here present UELer, an interactive viewer that links multi-channel image views with quantitative analysis results directly within Jupyter notebooks, requiring no dedicated infrastructure beyond the notebook session. Cells selected through computational analysis and summary plots can be inspected directly in their tissue context, and selections made in the image can be made available to any downstream analysis. Together, these capabilities support interactive data exploration, iterative cell annotation, and reproducible retrieval of selected regions.

**Availability and Implementation:** UELer is a Python package built on ipywidgets and runs in Jupyter environments supporting ipywidgets 8.1 or later, tested in JupyterLab and Visual Studio Code on Linux, macOS, and Windows. It is freely available under BSD 3-Clause license and can be installed via pip. Source code and documentation are available at https://github.com/HartmannLab/UELer and https://hartmannlab.github.io/UELer/. An online, no-install version runs remotely via BinderHub (https://mybinder.org/v2/gh/HartmannLab/UELer/main), accessible through the script/run_ueler_binder.ipynb notebook.

## 1. Introduction

Multiplexed imaging technologies are generating increasingly large and complex spatial datasets. As a result, computational analysis has become essential for extracting quantitative information from these images, and many analysis workflows are implemented in Python-based environments (Palla *et al*. 2022, Schapiro *et al*. 2022, Nirmal and Sorger 2024, Tan *et al*. 2025), commonly through Jupyter notebooks (Granger and Pérez 2021). Visual inspection remains critical, however, since quantitative features alone can miss biologically meaningful spatial patterns and fail to reveal common failure modes such as staining artifacts, segmentation errors, or ambiguous phenotypes.

Spatial proteomics datasets now often span dozens to hundreds of fields of view (FOVs) (Jackson *et al*. 2020, Schürch *et al*. 2020, Kinkhabwala *et al*. 2022, Najem *et al*. 2025, Vulliard *et al*. 2025, Eling *et al*. 2025, Liu *et al*. 2026, Lekiashvili *et al*. 2026), with tens to more than one hundred channels per image (Keren *et al*. 2019, Donovan *et al*. 2024, Kuehl *et al*. 2025), which complicates navigation, rendering, and interpretation. To support exploration of such multiplexed imaging data, a range of dedicated viewers has been developed. histoCAT couples single-cell gating to image views (Schapiro *et al*. 2017), Mantis Viewer adds segmentation-aware inspection of many channels (Schiemann *et al*. 2020). TissUUmaps brings performant web-based rendering (Pielawski *et al*. 2023), Odon optimizes rendering for whole-slide data (Coulton and McGranahan 2026), and CellTune targets iterative cell classification (Bussi *et al*. 2025). In parallel, napari (Chiu, Clack, and the napari community 2022) has become a foundation for image analysis in Python and supports spatial omics through plugins such as napari-spatialdata (Marconato *et al*. 2025). Vitessce provides coordinated views purpose-built for multimodal and spatially resolved single-cell data (Keller *et al*. 2025).

Despite this growing toolbox, practical deployment constraints still regularly force quantitative analysis and image inspection in separate environments. These tools are delivered either as desktop applications, which require a local display, or as browser-based applications, which require an additional data server even when embedded in a notebook. Satisfying these requirements is often non-trivial, especially on shared servers and compute clusters where large multiplexed imaging datasets are commonly analyzed. Retrieving spatial context for cells identified in computational analyses, or propagating image-based selections back to analytical results, therefore largely still requires manual export, import, and cross-referencing whenever these tools cannot be deployed alongside the analysis. This fragmentation is particularly limiting during manual data curation, where cell annotation requires repeated inspection of both quantitative summaries and spatial image context. Similarly, the identification and validation of larger tissue structures and multicellular neighborhoods benefits from interactive visualization of single-cell information and spatial context.

To address these limitations, we developed UELer (**U**nified **E**xploratory **L**inked View**er**), an interactive viewer that links multiplexed imaging data with quantitative analysis results within Jupyter notebook workflows. UELer communicates with the kernel over the connection the notebook already uses, requiring no additional server, port, or display. By enabling bidirectional navigation between image context and quantitative measurements, UELer brings exploration, validation and annotation into a single environment, without leaving the analysis workflow.

## 2. UELer: Linked exploration of multiplexed imaging data

UELer is implemented within the ipywidgets (https://github.com/jupyter-widgets/ipywidgets) ecosystem for use directly in Jupyter notebooks (Figure 1). It takes as input tabular single-cell measurements (CSV or AnnData (Virshup *et al*. 2024)) together with multiplexed image data, a multi-TIFF folder or a TIFF stack such as OME-TIFF (Goldberg *et al*. 2005), read either from local storage or directly from the BioImage Archive (Hartley *et al*. 2022). The interface is organized into three components: a viewer for image rendering and image-based interactions, a global settings system that maintains consistent rendering parameters across images, and linked plots that operate on summarized cell- and group-level measurements.

**Figure 1.**
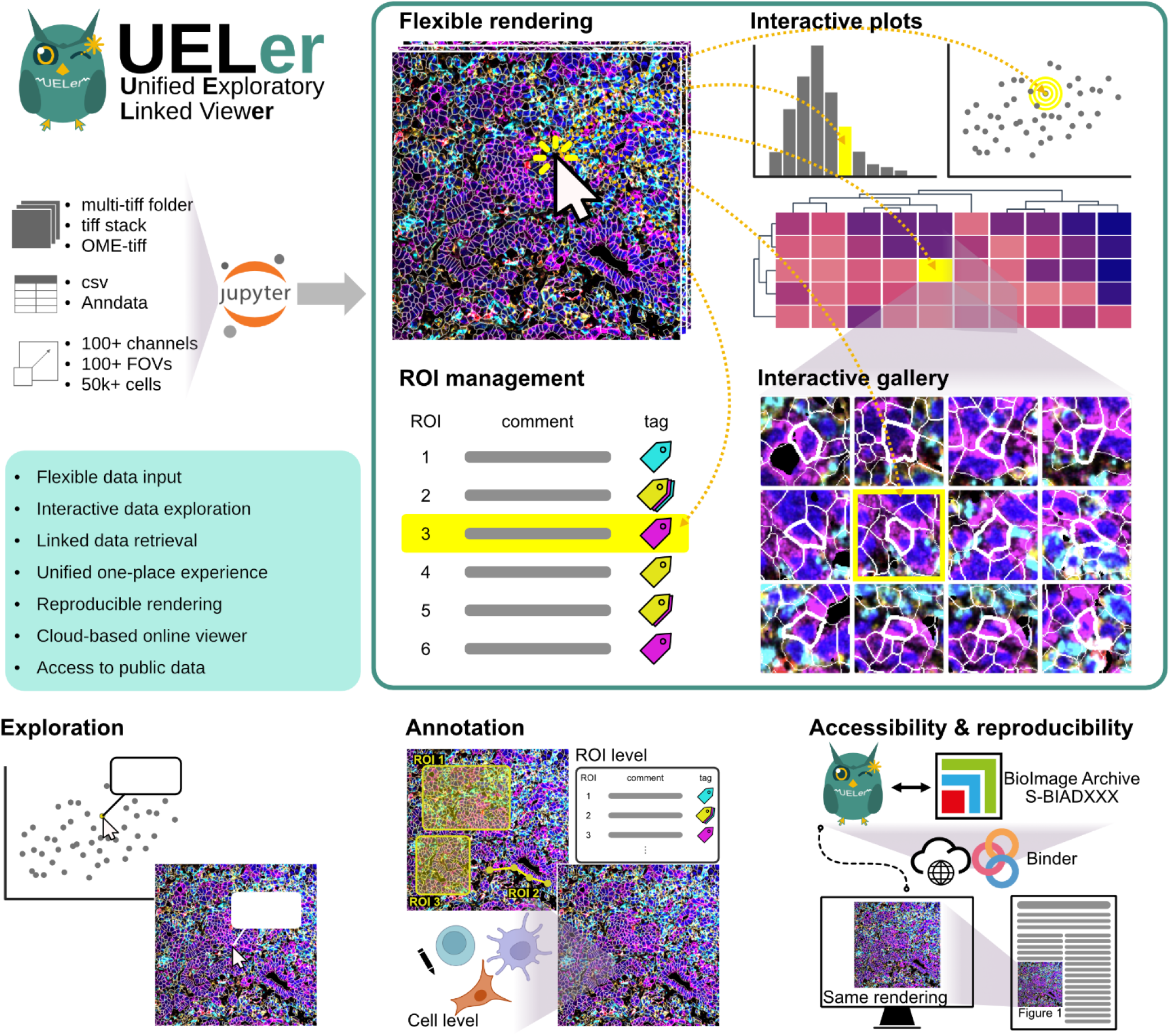
UELer links multiplexed image data with quantitative analysis in a single Jupyter notebook environment. UELer integrates tabular single-cell measurements (e.g., CSV or AnnData (Virshup *et al*. 2024)) with multiplexed image data (e.g., multi-TIFF folder or tiff stack, such as OME-TIFF (Goldberg *et al*. 2005)), stored locally or deposited on the BioImage Archive (Hartley *et al*. 2022), to enable linked exploration of spatial and quantitative information. The viewer combines multi-channel image rendering with interactive plots, cell galleries, and ROI management, allowing selections made in plots, images, or galleries to propagate across views.

The central interaction model connects multiplexed images and quantitative summaries through linked views. Cells shown in plots such as scatter plots, histograms, or dimensionality-reduction embeddings remain linked to their spatial coordinates in the tissue image through unique identifiers, so cells chosen in a plot can be inspected directly in the image. Importantly, selections propagate in both directions: any selection made in the image that can be expressed as a label, for example a gate, a phenotype, or a region assignment, is appended as a new column to the underlying single-cell data table (e.g. the AnnData object). The selection can be written back to the original data, thus making it immediately available to any downstream analysis in the notebook.

Image data are loaded lazily using Dask (Rocklin 2015), and rendering adapts to zoom level through downsampling. These mechanisms enable fast, responsive viewing of multiplexed imaging datasets spanning many channels and fields of view. Large viewports spanning multiple FOVs are rendered with a tiled approach that does not require pre-stitched images, ensuring that only the pixels in the current view are loaded into memory. Because the kernel runs alongside the data, only the rendered view is transmitted to the browser, and the volume crossing the notebook connection is independent of the size of the underlying dataset. Displaying five channels across a stitched view of approximately 100 FOVs (each 1024 × 1024 pixels at native resolution) completes in a few seconds at full zoom-out, where only a correspondingly downsampled subset of pixels is read.

Lastly, a plugin architecture allows custom analysis or visualization modules to be added to the viewer and its linked plots without modifying the core code, so that new components participate in linked exploration. The core plugin collection already spans quantitative display, visual inspection, curation, region management, and in-viewer analysis. Linked summary plots present measurements alongside the image (Scatter Plot, Histogram, Heatmap), whereas any per-cell value, discrete or continuous, can be rendered in spatial image context (Mask Painter) or as image crops of a selected subset (Cell Gallery). Curation plugins keep a version history across successive rounds of refinement (Cell Annotation) and attach tags to individual cells (Cell Table Editor). Spatial regions can be defined, tagged, and exported (ROI Manager, Batch Export), and clustering runs without leaving the viewer (FlowSOM).

## 3. Use Cases

### Interactive exploration and data retrieval

Exploration of multiplexed imaging datasets often begins with quantitative summaries, such as scatter plots or low-dimensional embeddings (e.g., UMAP). Although useful for identifying patterns and cellular populations, these representations abstract away spatial context, making it difficult to interpret cellular phenotypes or validate computational stratifications. Visual inspection of the corresponding cells and tissue regions is therefore often necessary but retrieving the specific images that carry the cells of interest can require multiple manual steps.

Using UELer, cells selected in data analysis summaries can be inspected immediately within their original tissue context or displayed in the Cell Gallery plugin. This allows users to evaluate cellular morphology, the local microenvironment, staining quality, and segmentation accuracy. Moreover, plugins are interconnected, enabling selections to be examined using the visualization tools best suited to each analytical task.

For example, in (Liu *et al*. 2026, Lekiashvili *et al*. 2026), we used the Scatter Plot plugin to identify cells that appeared positive for both SMA, a stromal marker, and CD45, an immune marker. We selected this population in the scatter plot using the lasso tool, which prompted the Cell Gallery plugin to display images of selected cells in batches of 20. Visual inspection of these images showed that many apparent double-positive events were not genuinely co-expressing cells, but instead represented closely interacting fibroblasts and immune cells with spillover or fibroblast–immune-cell doublets. Clicking on individual images in the Cell Gallery plugin navigated directly to the corresponding *in situ* locations in the main viewer. This spatial inspection further showed that these events were concentrated in dense stromal regions containing abundant immune cells and fibroblasts, providing a spatial explanation for the apparent double-positive population.

Together, this illustrates that connecting a quantitative pattern to its spatial context distinguishes biological phenotypes from segmentation artifacts that are not identifiable from single-cell data alone.

### Iterative cell annotation workflows

Manual annotation of cell populations often requires repeated transitions between computational analysis and visual inspection. While automated clustering methods can identify candidate populations, heterogeneous or poorly separated clusters cannot always be assigned confidently from marker-expression profiles alone and therefore require iterative examination and refinement.

A typical workflow supported by UELer begins with unsupervised clustering based on selected lineage markers, followed by the aggregation of clusters into meta-clusters according to their marker-expression profiles and hierarchical relationships. Ambiguous clusters can then be examined at the single-cell level and re-clustered as needed until biologically coherent populations emerge. This same strategy can subsequently be applied within individual meta-clusters to resolve finer cell subtypes.

For example, in (Liu *et al*. 2026, Lekiashvili *et al*. 2026), we partitioned over 550,000 cells into 100 clusters using FlowSOM through UELer’s FlowSOM plugin. We visualized cluster-level marker profiles (including epithelial, hepatocyte, stromal, and immune markers) with the Heatmap plugin and generated initial meta-clusters by cutting the associated hierarchical clustering tree. Clusters with ambiguous identities, for example, those showing comparable abundance of markers associated with multiple lineages, were evaluated by reviewing images of dozens to hundreds of cells sampled across all FOVs using the Cell Gallery plugin. These clusters were then iteratively re-clustered, inspected, and refined. Finally, biologically interpretable labels (e.g., cancer cells, immune cells, and fibroblasts) were assigned to the resulting meta-clusters in the Heatmap plugin and written back to the cell table for downstream analysis.

In summary, by keeping computational groupings connected to their visual evidence throughout refinement, UELer supports cell annotation as an iterative process of biological interpretation rather than a sequence of disconnected analytical and manual steps.

### Reproducible exploration and dataset sharing

Representative tissue regions and single-cell views are frequently included in scientific presentations and publications. However, revisiting these same views can be difficult without dedicated ROI management. Reproducing an identical visualization presents an additional challenge when viewers store only ROI geometry without the associated rendering parameters, such as selected markers, colors, intensity scales, or mask settings. Consequently, previously generated images may be difficult to reproduce or share consistently with collaborators and audiences.

UELer’s ROI manager and records allow users to restore the same spatial view and rendering configuration across sessions, supporting reproducible inspection, figure generation, and collaboration. UELer’s plugin ecosystem further extends the ROI Manager beyond a storage tool by connecting saved views to downstream visualization and export workflows.

For example, to visualize MIBI data for (Liu *et al*. 2026, Lekiashvili *et al*. 2026), we first defined marker sets specifying the markers, their rendering colors, and their intensity scales. We then navigated to a representative region in the main viewer and captured the current view using the ROI Manager. Each saved ROI was associated with a marker set, a mask type, and user annotations in the form of free-text comments and tags, such as “Figure 1”, “stroma”, and “immune infiltration”. Saved ROIs could then be filtered using these tags, allowing relevant views to be retrieved quickly. Collecting multiple ROIs, we used the Batch Export plugin to export the corresponding visualizations as PDF files, which were subsequently combined with other figure components in a graphics-layout application. Because the rendering parameters were stored with each ROI, the exported views could be reproduced without manual reconstruction.

These capabilities extend reproducibility from figure generation to dataset sharing because the BinderHub link in the UELer repository launches a session in the browser without local installation. This enables interactive exploration of published multiplexed imaging datasets hosted in repositories such as the BioImage Archive (Hartley *et al*. 2022). When saved ROIs accompany a dataset user can revisit representative regions and inspect the underlying spatial data before downloading the complete dataset.

In this way, representative images become reproducible entry points into published datasets rather than static illustrations.

All the use cases were run remotely on the compute cluster that hosts the data and the analysis. Meanwhile, cross-platform compatibility allowed co-authors to reproduce the rendering on their own devices.

## 4. Conclusions

Multiplexed imaging datasets increasingly require workflows that combine computational analysis with careful visual inspection of spatial context. By linking quantitative analysis results to multiplexed image views within Jupyter notebooks, and by writing image-based selections back into the analysis object, UELer keeps exploration, validation, and annotation inside a unified workflow that runs wherever the notebook itself runs. UELer thus complements existing visualization platforms by focusing on tight integration with Python analysis pipelines. Keeping analysis code, data, and interactive visualization in a single document directly supports more transparent and reproducible exploration of multiplexed imaging datasets.

## Acknowledgements

We thank Amirhossein Sohrabipour and Vu Ha Anh Ta for testing the software and providing feedback, and Loan Vulliard and Miray Cetin for scientific input. The authors used Claude models Opus 4.6 and 5 within Claude Code for code assistance during development of UELer and with the chat mode for language editing of this manuscript. All output was reviewed and verified by the authors, who wrote the manuscript and take full responsibility for its content and for the correctness of the software.

## Author contributions

Yu-Le Wu: Conceptualization (lead); software (lead); validation (lead); visualization (lead); data curation (lead); writing – original draft (lead); writing – review and editing (lead). Chun-Shan Liu: Resources (lead); data curation (supporting); validation (supporting); writing – review and editing (supporting). Bénédicte Lenoir: Resources (supporting); data curation (supporting). Kilian Merz: Resources (supporting); data curation (supporting); writing – review and editing (supporting). Michael T. Dill: Supervision (supporting); funding acquisition (supporting). Felix J. Hartmann: Supervision (lead); funding acquisition (lead); writing – review and editing (supporting).

## Conflict of Interest

The authors declare no conflict of interest.

## Funding

F.J.H. is supported by the Initiative and Networking Fund of the Helmholtz Association under the Helmholtz Young Investigator Program (VH-NG-1605), the State Parliament of Baden-Württemberg for the Innovation Campus Health + Life Science Alliance Heidelberg Mannheim, the Rolf M. Schwiete Foundation, and the Hector Foundation II. Funded by the Deutsche Forschungsgemeinschaft (DFG, German Research Foundation) under Germany’s Excellence Strategy – EXC 3018/1 – 533587280. Funded by the European Union (ERC, SpatialTMEMetabolism, 101116823). Views and opinions expressed are however those of the author(s) only and do not necessarily reflect those of the European Union or the European Research Council Executive Agency. Neither the European Union nor the granting authority can be held responsible for them. M.T.D is supported by the Rolf M. Schwiete Stiftung and a Max Eder grant (70113858) from the German Cancer Aid. Y.-L.W. is supported by a DKFZ Postdoctoral fellowship.

## Notes

https://github.com/HartmannLab/UELer

https://hartmannlab.github.io/UELer/

### Competing Interest Statement

The authors have declared no competing interest.

